# Contribution of common and rare variants to bipolar disorder susceptibility in extended pedigrees from population isolates

**DOI:** 10.1101/363267

**Authors:** Jae Hoon Sul, Susan K. Service, Alden Y. Huang, Vasily Ramensky, Sun-Goo Hwang, Terri M. Teshiba, YoungJun Park, Anil P. S. Ori, Zhongyang Zhang, Niamh Mullins, Loes M. Olde Loohuis, Scott C. Fears, Carmen Araya, Xinia Araya, Mitzi Spesny, Julio Bejarano, Margarita Ramirez, Gabriel Castrillón, Juliana Gomez-Makhinson, Maria C. Lopez, Gabriel Montoya, Claudia P. Montoya, Ileana Aldana, Javier I. Escobar, Jorge Ospina-Duque, Barbara Kremeyer, Gabriel Bedoya, Andres Ruiz-Linares, Rita M. Cantor, Julio Molina, Giovanni Coppola, Roel A. Ophoff, Gabriel Macaya, Carlos Lopez-Jaramillo, Victor Reus, Carrie E. Bearden, Chiara Sabatti, Nelson B. Freimer

## Abstract

Current evidence from case/control studies indicates that genetic risk for psychiatric disorders derives primarily from numerous common variants, each with a small phenotypic impact. The literature describing apparent segregation of bipolar disorder (BP) in numerous multigenerational pedigrees suggests that, in such families, large-effect inherited variants might play a greater role. To evaluate this hypothesis, we conducted genetic analyses in 26 Colombian (CO) and Costa Rican (CR) pedigrees ascertained for BP1, the most severe and heritable form of BP. In these pedigrees, we performed microarray SNP genotyping of 856 individuals and high-coverage whole-genome sequencing of 454 individuals. Compared to their unaffected relatives, BP1 individuals had higher polygenic risk scores estimated from SNPs associated with BP discovered in independent genome-wide association studies, and also displayed a higher burden of rare deleterious single nucleotide variants (SNVs) and rare copy number variants (CNVs) in genes likely to be relevant to BP1. Parametric and non-parametric linkage analyses identified 15 BP1 linkage peaks, encompassing about 100 genes, although we observed no significant segregation pattern for any particular rare SNVs and CNVs. These results suggest that even in extended pedigrees, genetic risk for BP appears to derive mainly from small to moderate effect rare and common variants.

## Introduction

Bipolar disorder (BP), consisting of episodes of mania and depression, has a heritability from twin studies estimated to be about 80%(1). For BP, as for most other common disorders, SNP-based genome-wide association studies (GWAS) of large case/control samples have discovered many loci that contribute unequivocally to disease risk but that collectively explain only a small fraction of disease heritability. The most recent published BP GWAS, incorporating more than 20,000 cases and 30,000 controls, has reported 30 genome-wide significant SNP-associations and SNP-based heritability (h^2^_snp_) of 25% for BP1(2). The hypothesis that rare single nucleotide variants (SNVs) and rare copy number variants (CNVs) could explain a substantial proportion of this “missing heritability”(3) has motivated the rapid growth of whole exome sequencing (WES) and whole genome sequencing (WGS) throughout biomedicine, including psychiatry. More than for other psychiatric disorders, however, sequencing efforts to identify variants with a high impact on BP risk have continued to focus on pedigrees(4-7). This focus reflects published descriptions, over several decades, of numerous extended families in which BP is observed across multiple generations; as would be expected if these pedigrees were segregating a relatively high-penetrance susceptibility variant.

Because the literature regarding the apparent segregation of BP in extended pedigrees is mostly anecdotal(7-9), we aimed to systematically characterize the genetic contribution to BP disease risk in a series of such families through evaluation of variants across the allele frequency spectrum. If rare variants contribute to this risk it is expected that they would be enriched in this sample which is, to our knowledge, the largest BP pedigree sample sequenced to date. Additionally, because a wide range of evidence indicates considerable etiological heterogeneity between BP1 and milder forms of BP, this study focused exclusively on families ascertained for multiple cases of BP1, a strategy which we reasoned would reduce the impact of such heterogeneity. In a further effort to reduce heterogeneity, we limited the dataset to pedigrees derived from two Latin American populations that are considered closely related genetic isolates; the province of Antioquia in CO and the Central Valley of CR(10).

We collected microarray SNP data for 856 family members (as reported previously)(11), and performed high-coverage whole-genome sequencing (WGS) on 454 individuals, selected because their position in the pedigrees indicated that they would provide substantial information for imputing rare variation in the remaining family members. We analyzed these data to obtain high-quality genotypes for several categories of genetic variant, including SNVs, CNVs, short tandem repeats (STRs), and *de novo* mutations (DNMs). With this rich genetic information, we sought to evaluate the impact of both common and rare variants on BP1, focusing on two major questions about the genetic etiology of BP1. The first question relates to the overall genetic architecture of BP1 in these families. We attempted to address this question by characterizing the overall contribution of genetic variation to BP1 using both common and rare variants in a genome-wide burden analysis. For common variants, we calculated the polygenic risk scores (PRS) using the latest BP1 GWAS summary statistics(2) and compared the polygenic burden of risk alleles in affected cases and related controls. For rare variants, the genome-wide burden analysis contrasted the burden of rare deleterious variants in a set of genes related to BP1 between affected cases and related controls, where deleterious variants are defined as stop-gain, stop-loss, splice-site, or missense variants predicted to be damaging by Polyphen-2(12). The second question we focused on in this study is identifying specific genetic loci contributing to BP1 or segregating in the families (segregation analysis). To address this question, we performed linkage analysis to test segregation of common variants and employed a new method to test segregation of rare deleterious variants in families.

## Results

### Genotyping and sequencing of CO/CR families ascertained for BP1

Our study included 26 extended families (11 CO and 15 CR) whose pedigree size varied from 12 to 355 individuals, with a range of 2 to 44 known BP1 individuals per pedigree (Table 1). Methods used to assign diagnoses of BP1 in these pedigrees have been described previously(13) and are outlined briefly in the Methods section of this paper, along with the criteria used to designate relatives as controls. We have performed SNP genotyping on 856 individuals; details of SNP genotyping of these pedigrees, its QC, and its application to analyses of sleep and circadian phenotypes have been reported previously(11). After QC, we have 838 individuals including 206 BP1 individuals genotyped at 2,026,257 SNPs (Figure S1). Here we use these genotype data for BP1 linkage analysis and for estimation of inheritance vectors (IVs) for genotype imputation of WGS data.

**Table 1.** Description of families included in the current study. FamID: Family ID; N: number of individuals in the family; NBP: number of BP1 individuals in the family; NGeno: number of genotyped family members; NPheno: number of family members with endophenotype data; NMale: number of males in the family.

| FamID | N | NBP | NGeno | NPheno | NMale |
| --- | --- | --- | --- | --- | --- |
| CO10 | 38 | 6 | 24 | 24 | 13 |
| CO13 | 24 | 5 | 20 | 19 | 10 |
| CO14 | 29 | 8 | 23 | 22 | 16 |
| CO15 | 27 | 5 | 21 | 21 | 12 |
| CO18 | 37 | 6 | 26 | 25 | 18 |
| CO23 | 48 | 9 | 32 | 31 | 20 |
| CO25 | 15 | 4 | 13 | 12 | 7 |
| CO27 | 58 | 9 | 35 | 35 | 31 |
| CO4 | 73 | 10 | 42 | 43 | 37 |
| CO7 | 149 | 29 | 111 | 112 | 72 |
| CO8 | 16 | 5 | 7 | 8 | 6 |
| CR001 | 46 | 8 | 20 | 7 | 25 |
| CR004 | 187 | 23 | 77 | 45 | 91 |
| CR006 | 37 | 4 | 13 | 8 | 22 |
| CR007 | 12 | 2 | 9 | 6 | 7 |
| CR008 | 30 | 7 | 17 | 13 | 15 |
| CR009 | 44 | 9 | 32 | 34 | 17 |
| CR010 | 30 | 4 | 17 | 12 | 15 |
| CR011 | 16 | 3 | 13 | 12 | 6 |
| CR012 | 35 | 5 | 26 | 22 | 16 |
| CR013 | 39 | 4 | 10 | 8 | 15 |
| CR014 | 26 | 5 | 8 | 3 | 14 |
| CR015 | 19 | 2 | 10 | 10 | 7 |
| CR016 | 24 | 4 | 18 | 19 | 14 |
| CR201 | 355 | 44 | 201 | 177 | 176 |
| CR277 | 25 | 4 | 13 | 10 | 11 |

We have performed high-coverage (36x) WGS on 454 individuals in 22 families (10 CO and 12 CR). SNVs were called following the best practice pipeline of GATK HaplotypeCaller(14), and variant-level QC was performed using a classifier that employs sequencing quality information to filter out poorly sequenced variants, resulting in 20,396,290 autosomal SNVs (Table S1, see Methods). Genotype calls were then refined with Polymutt(15), a pedigree-aware variant calling algorithm that corrected 99.83% of Mendelian inconsistent genotypes by changing mostly low quality genotype calls (Table S2). We verified the ethnicity of founders using principal component analysis (PCA) with 1000 Genomes(16) (Figure S2). After removal of WGS data from five individuals that failed QC (Figure S1), we used the WGS data from the remaining 449 directly sequenced individuals to impute genome-wide genotypes in 334 individuals for whom we had only microarray data. We performed pedigree-aware imputation using the GIGI software(17), and measured imputation accuracy by comparing genotype concordance between microarray and imputed data for each individual. We found that, except for one individual, who we later excluded, the individuals whose genotypes we imputed displayed a genotype concordance rate > 97.93% between the microarray data and the GIGI imputation data (Figure S3). After imputation and QC, our WGS dataset consisted of 782 individuals including 190 BP1; 130 unaffected individuals, who have passed the age of risk and whom we considered as controls; and 462 individuals with unknown disease status. We used the BP1 and control data to perform genome-wide burden analyses of common and rare variants and segregation analyses of rare variants.

We also used the WGS data to call STRs in the dataset, using the lobSTR software(18). We devised a filtering strategy to extract high-quality STR calls by comparing the lobSTR STR calls to previously obtained STR data identified with electrophoresis, in one of the pedigrees (Table S3). After QC filtering, we detected 86,601 STR calls which we used in linkage analysis. We also performed genome-wide detection of CNVs using both microarray and WGS data. We applied a previously established pipeline for microarray-based detection(19). Of 838 individuals passing SNP QC, 782 (93.3%, 189 BP1 individuals, 128 controls, and 465 unknown) also passed QC based on intensity metrics and overall CNV load (Figure S4). Note that these 782 individuals are not the same set of individuals as the aforementioned 782 individuals after GIGI imputation because the number of BP1 individuals and controls differs between the two due to different QC and different number of families analyzed (26 for microarray data vs. 22 for WGS data). We detected a total of 5,437 CNVs (3,317 deletions and 2,120 duplications) after filtering for rare events > 5kb in length and spanned by a minimum of 10 probes (see Methods). To detect CNVs from the WGS data, we used Genome STRiP software (20, 21) to discover and genotype only bi-allelic deletions, as it has been demonstrated that such variants are far more amenable to imputation than other structural variant classes(21). After performing QC and merging of redundant sites, we genotyped 8,768 distinct deletion loci across the 449 individuals passing both SNV and CNV-based QC, and used these calls to impute CNVs in the same set of individuals imputed for SNVs. Lastly, we detected DNMs among 67 sequenced trios using the TrioDenovo software(22). Detailed results about quality of variant calling and QC are discussed in Supplementary Text. A summary on the number of variants and individuals after QC for different types of variants, and on which analysis is applied to each type of variant, is discussed in Table S4.

### Characteristics of admixture in the CO/CR Extended Pedigrees

Individuals in our CO/CR pedigrees were from two Latin American populations that mainly derive from African, European, and Native American ancestries(23). We estimated genome-wide ancestry proportions in members of the CR and CO pedigrees using ADMIXTURE(24)(v1.22). We found that the majority of ancestry was European, while the Native American proportion was also substantial (Figure S5). We also discovered that the admixture proportions in these pedigrees were significantly related to BP1 status; risk of BP1 increased by odds ratio (OR) of 1.53 (p=0.0008) with every increase of 0.1 units of European ancestry, while we observed the opposite trend for Native American ancestry with OR of 0.67 (p=0.0096), and African ancestry with OR of 0.61 (p=0.026) (Figure S6).

### The Genetic Architecture of BP1 in the CO/CR Extended Pedigrees

We evaluated the contribution of rare and common variants to the risk of BP1 through two genome-wide burden analyses comparing the BP1 individuals to the controls. To evaluate common variant genome-wide burden we computed the PRS using independent GWAS results for BP1 and SCZ. To evaluate rare variant genome-wide burden we estimated the mean number of alternative alleles at loci thought to be important to BP1. All analyses accounted for relationships among family members using theoretical kinship and global estimates of European ancestry using mixed models. Results using kinship derived from genetic data were not substantively different (data not shown).

#### PRS analysis of BP1 and SCZ GWAS summary statistics

To determine the effect of common SNPs on BP1 in the CO/CR pedigrees, we calculated a PRS for each individual using the latest Psychiatric Genomics Consortium (PGC) GWAS summary statistics for BP1, consisting of 14,583 cases and 30,424 controls(2). A higher PRS indicates a greater polygenic burden of risk alleles for BP1. We calculated the PRS at different GWAS p-value thresholds, where higher p-value thresholds used more common variants in the PRS calculation than did lower p-value thresholds. Results show that the mean PRS is higher in the 190 BP1 individuals compared to 130 controls, at GWAS p-value thresholds of 0.01 and 0.001 using a linear mixed model (LMM) (p = 0.001 and 0.007, respectively) and at a GWAS p-value threshold of 0.01 using a generalized linear mixed model (GLMM) (p=0.003, Table 2, Figure 1). We also calculated Nagelkerke’s R^2^ from logistic regression and found that these PRS explain 1.5% of the variance (Table S5). This R^2^ is noticeably smaller than that explained by PRS in the latest PGC BP GWAS data where the weighted average Nagelkerke’s R^2^ is 8%. We considered the possibility that this difference in the variance explained by the PRS might reflect population-level differences between the mostly European-descended PGC samples, and the Latin America pedigrees in our study, which had a substantial component of Native American and African ancestry. However, as more than 90% of the SNPs in the PGC BP GWAS were present in the CO/CR pedigrees, we consider it unlikely that this explanation, alone, explains the difference between the pedigree and population samples (Table S5).

**Table 2.** Comparison of Polygenic risk score estimated from PGC BP1 GW AS summary statistic between BP1 individuals and controls. P-values are computed using linear and logistic regression models by taking into account relatedness; BP: coefficients and p-values for BP1 status; AdMix: coefficients and p-values for global admixture proportions of European ancestry; LM: Linear model; LR: Logistic regression; QNPRS: quantile-normalized polygenic risk scores.

| GWAS Threshold | NSNPs | LM Beta (BP) | LM P-value (BP) | LM Beta (AdMix) | LM P-value (AdMix) | LR Log OR (QNPRS) | LR P-value (QNPRS) | OR for 1-unit increase in QNPRS | LR Log OR (AdMix) | LR P-value (AdMix) |
| --- | --- | --- | --- | --- | --- | --- | --- | --- | --- | --- |
| 0.01 | 26868 | 0.27 | 0.0014 | -4.11 | 2.13E-09 | 0.39 | 0.0027 | 1.477 | 5.53 | 5.12E-05 |
| 0.001 | 4241 | 0.25 | 0.0069 | -3.24 | 2.92E-05 | 0.23 | 0.0539 | 1.262 | 4.90 | 2.27E-04 |
| 0.0001 | 789 | 0.06 | 0.4842 | -2.83 | 1.23E-04 | 0.13 | 0.2704 | 1.143 | 4.54 | 4.52E-04 |
| 0.000001 | 65 | 0.09 | 0.3421 | -0.44 | 5.81E-01 | 0.03 | 0.7740 | 1.034 | 4.29 | 7.48E-04 |
| 0.00000005 | 13 | 0.05 | 0.6136 | -0.74 | 3.67E-01 | 0.00 | 0.9709 | 0.996 | 4.27 | 7.94E-04 |

**Figure 1.**
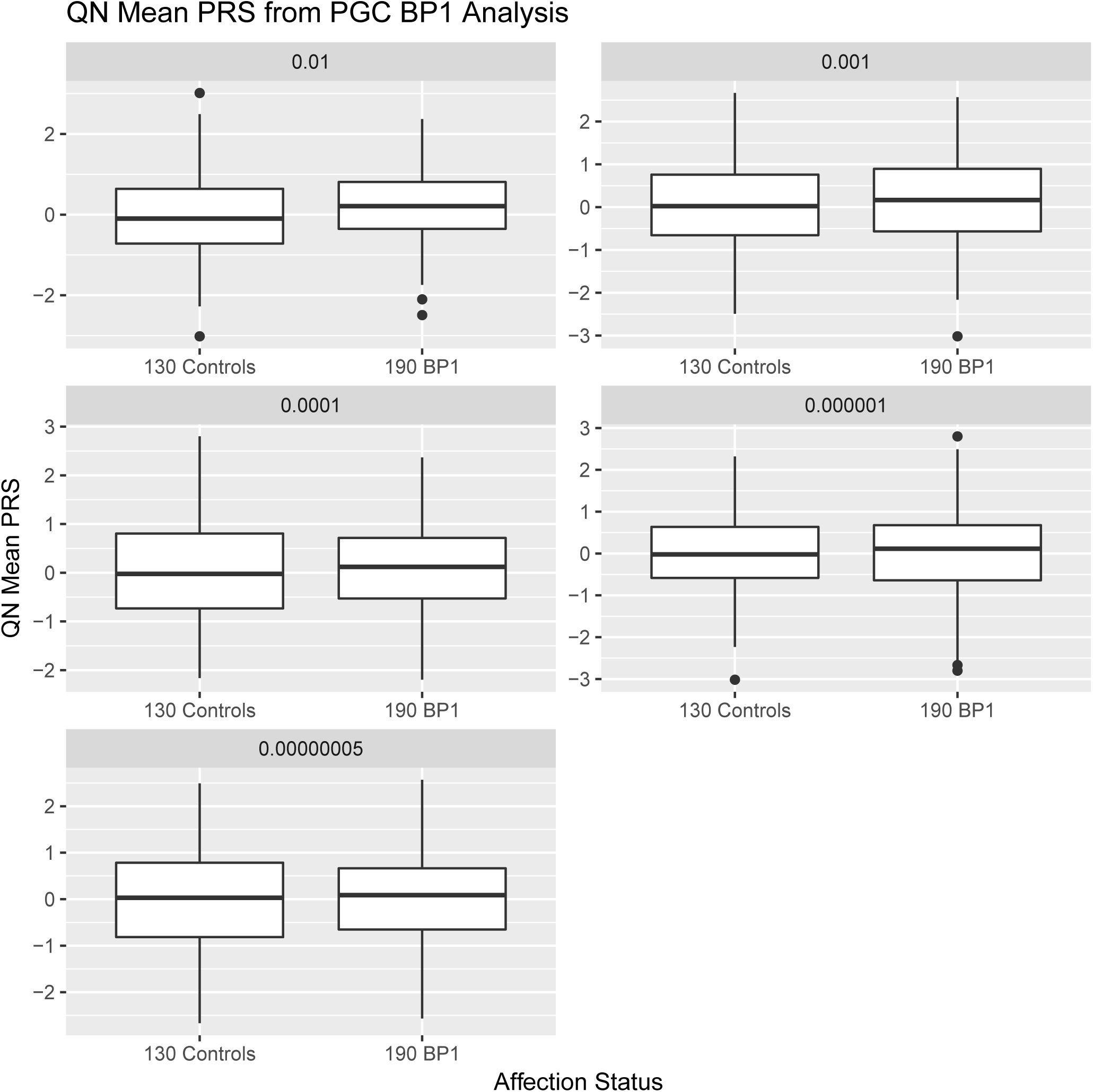
Comparison of PRS estimated from PGC BP1 GWAS summary statistics in BP1 individuals and controls. PRS is computed at different GWAS p-value thresholds of the PGC BP1 GWAS.

Although evidence over several decades delineated the distinctions between BP and SCZ, more recent studies have highlighted genetic overlaps between these syndromes(2, 25, 26), which share symptoms in common. Notably, in GWAS data from large BP case/control samples, the PRS estimated from the PGC’s SCZ GWAS results have explained up to 2.5% of BP variance(25). We contrasted the mean PRS from SCZ GWAS in BP1 affected individuals and related controls in the 22 pedigrees in our study for which we had WGS data; to calculate the PRS we used the latest available GWAS summary statistics for schizophrenia(27), derived from 36,989 cases and 113,075 controls. These SCZ PRS are not associated with an increased risk of BP1, in the CO/CR pedigrees, at any of the GWAS p-value thresholds that we examined (Table S6, Figure S7). The association of SCZ PRS with BP1 in the PGC, but not in the CO/CR pedigrees may suggest differences in the characteristics of BP1 between these samples; in particular, this contrast between our results and those of the PGC may reflect the fact that we ascertained each of the pedigrees for multiple closely related cases of BP1.

#### Identification of a gene-set implicated in BP1

To increase the statistical power to detect the effect of rare variants on BP1, we sought to limit the number of such variants to be evaluated by focusing on genes for which *a priori* information indicated their relevance to the phenotype. To identify such genes, we utilized three sources of information: (1) functional annotations of genome regions that have demonstrated increased heritability for BP1 in PGC GWAS data; (2) a list of the genes in the vicinity of genome-wide significant BP1 associations in the most recent PGC GWAS of BP; (3) a list of the genes located under linkage peaks for BP1 in our analyses of these pedigrees. For each source, we identified genes that contain at least one putatively deleterious SNV in our dataset.

First, we evaluated the latest PGC BP1 GWAS summary statistics(2) to identify cell-type specific promotor or enhancer regions in which BP1 heritability is enriched compared to the genome overall, using the methods and cell-type groups employed by Finucane et al. in a study of an earlier PGC GWAS with a broader definition of BP(28). Consistent with their results, among the 10 cell-type groups that they had identified, we observed significant enrichment of heritability for BP1 only in the central nervous system (CNS) cell type group (Figure S8). The CNS cell type region comprised 457.5Mb of the genome, and contained 8,714 genes with at least one deleterious SNV in our dataset.

Second, from clumping the PGC BP1 GWAS results using LD evaluated in founders of the CO/CR pedigrees, we identified a total of 15 genome-wide significant lead SNPs, with 72 genes in the 6.35Mb regions around these SNPs (see Methods). Finally, in evaluating the regions defined by peaks from the BP1 linkage analysis, which we discuss in more detail below, we identified 99 genes sited within 1Mb of these peaks. Ignoring overlap of genes appearing in multiple sources, the three sources yielded a gene-set of 8,757 unique genes for our rare variant analyses of BP1.

#### Burden of rare deleterious SNVs and CNVs in the gene-set for BP1

We defined rare SNVs using both an external source of allele frequency (Colombians in the 1000 Genomes database(16) and Latinos in the ExAC database(29)) and allele frequency observed in the CO/CR families. We used an MAF threshold of 1% for the external source and 10% for allele frequency estimated from the CO/CR families (see Methods). We identified 25,072 rare deleterious SNVs in 8,236 of the 8,757 genes in our gene-set. For each individual, we computed the mean genome-wide burden of these rare deleterious SNVs, then compared these means between BP1 individuals and related controls, while taking into account the proportion of European ancestry in each individual, and the mean genome-wide burden of all rare SNVs in the gene-set. The latter procedure is necessary to account for individual differences in total amount of variation; an individual is likely to carry more rare deleterious SNVs if she/he carries more rare SNVs overall. We found that the mean burden of the rare deleterious SNVs was higher in BP1 individuals than in controls (p=0.047 using LMM, Figure S9). The risk of BP1, as indicated by odds ratio (OR) increased by 1.26 for every one unit increase in quantile-normal transformed residual mean burden (p=0.048 using GLMM).

We also performed a burden analysis using rare CNVs. For CNVs called using microarray data, we restricted our analysis to variants greater than 5kb in length and spanned by a minimum of 10 probes, and used frequency information from the Database of Genomic Variants (30) to define a set of rare variants. We identified 2,186 rare CNVs among 189 BP1 individuals and 128 related controls, and observed that the average number of such CNVs genome-wide did not differ between BP1 individuals and controls (6.86 and 6.95, respectively; p=0.67 with OR of 0.99 using GLMM). For each sample, we measured the CNV burden by enumeration of the gene count, defined as the number of genes in our BP1 gene-set intersected by rare CNVs carried by that individual. We measured enrichment of the CNV burden in BP1 individuals, under both LMM and GLMM, accounting for factors known to affect global measures of genic CNV burden, including both the total number of CNVs and average CNV length (31), as well as genotyping batch. The BP1 individuals had a higher mean burden of genes in our BP1 gene-set affected by rare CNVs than controls (p=0.013 using LMM, Figure 2, Table S7), and this higher burden increased the risk of BP1 with OR of 1.34 for every one unit increase in quantile-normal transformed residual of the mean number of genes affected (p=0.018 using GLMM). Stratifying our analysis by CNV type, we found that this increased burden was attributable exclusively to deletions (p=2.2e-3 using LMM and p=3.8e-3 with OR of 1.44 using GLMM). We repeated this analysis for deletions generated by imputation of CNV calls detected from WGS, filtering our rare deletions based on structural variant calls from Phase 3 of the 1000 Genomes Project(32). Rare deletions detected from WGS data (n=4,436 in 320 samples consisting of 190 BP1 individuals and 130 controls) were far more numerous than those detected from a subset of this sample used in the microarray analysis (n=2,186 in 317 samples consisting of 189 BP1 individuals and 128 controls, a subset of the sample used in WGS data). It is important to note that the set of BP1 individuals and controls is different between WGS CNV data and microarray CNV data because QC and the number of families (22 for WGS CNV data and 26 for microarray CNV data) were different. Similarly, although there was no difference in the overall genome-wide rate of rare deletions between cases and controls (13.71 and 14.08, respectively; p=0.45 with OR of 0.98 using GLMM), we observed an increased burden of genes in the BP1 gene-set affected by rare CNVs from WGS for BP1 individuals (p=0.022 using LMM), which increased the risk of BP1 with OR of 1.3 for every one unit increase in quantile-normal transformed residual of the mean number of genes affected (p=0.033 using GLMM). This burden was greater (p=6.1e-3 using LMM and p=8.7e-3 with OR of 1.39 using GLMM) when restricting our analysis to the subset of CNVs covered by a minimum of 10 SNPs on the Omni2.5 array (n=1,511), thus demonstrating a consistent increase in gene count burden using different methods of detection.

**Figure 2.**
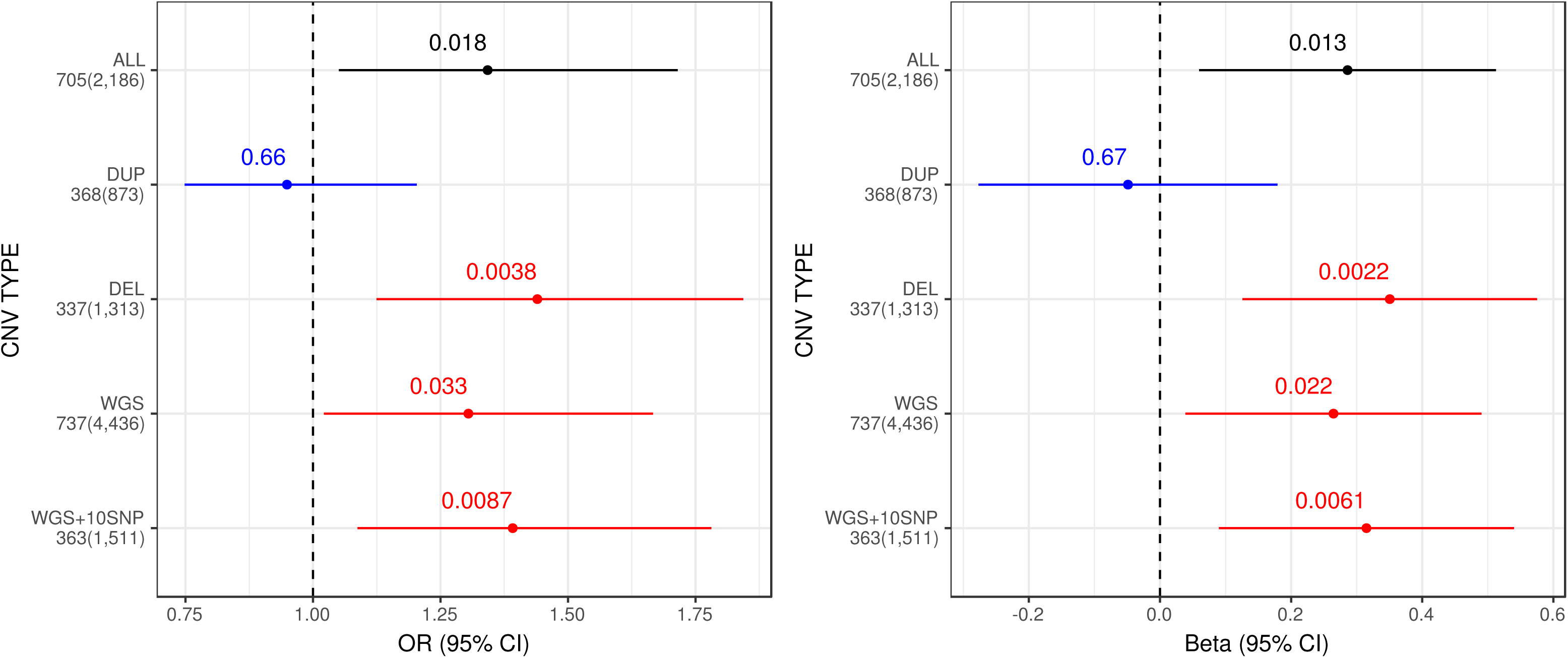
Forest plot depicts the increased burden of rare CNVs among BP1 individuals affecting BP1 related genes, stratified by a detection method (microarray or WGS) and a CNV type. The total number of CNVs detected and the number of CNVs affecting BP1 related genes are displayed for each category. To correct for individual relatedness and other potential confounders (see Methods), enrichment was assessed using a logistic mixed model (GLMM, left) and a linear mixed model (LMM, right).

### Genetic Loci Segregating with BP1 in the CO/CR Extended Pedigrees

#### Linkage analysis using microarray and STR data

To identify loci that segregate with BP1 in the CO/CR pedigrees, we performed linkage analysis using both common SNPs from microarray data (209 BP1 individuals in 26 families) and STRs from WGS data (143 BP1 in 22 families). For these analyses, we considered individuals diagnosed with BP1 as affected, and designated the phenotype of all other individuals as unknown. For the linkage analysis with common SNPs, we used 99,446 independent bi-allelic SNPs with MAF > 0.35, while for the analysis with STRs, we used 85,813 bi- and multi-allelic STRs. We applied two-point parametric linkage analysis, using the Mendel software(33), to both SNP and STR data and estimated the HLOD (LOD with heterogeneity) scores. While there is not a single established approach for assessing significance in this setting, we report here the locations of HLODs greater than 4.1, which correspond to the LOD threshold for genome-wide significant linkage of 3.3 suggested by Lander and Kruglyak(34) (see Methods). We also performed non-parametric linkage (NPL) analysis on these data using the Rapid software package(35), and estimated p-values via simulations.

Four microarray SNPs displayed HLODs exceeding 4.1 (Figure 3, Table 3); two SNPs on 2q33 and 2q36 (kgp5681631 at 206.8 Mb and kgp10402924 at 228.3 Mb, with HLODs of 4.71 and 4.50, respectively) and two SNPs on 16q23 (kgp2684721 at 79.5 Mb and rs8058020 at 83.7 Mb, with HLODs of 4.13 and 4.59, respectively). There was no heterogeneity for the linkage results on 2q33 and 2q36, and on 16q23 the estimate of the proportion of linked families (denoted as *a*) was 0.90. The HLOD for two STRs exceeded the threshold of 4.1, one on 1q44 (247.1 Mb, HLOD=5.03, *a* = 0.72), and one on 15q14 (35.3 Mb, HLOD=4.32, *a* = 0.62). While these results at five loci are suggestive, our simulation studies indicate that the evidence in favor of linkage is not as strong as it might appear. We carried out simulations under the null hypothesis of no linkage to BP1 anywhere in the genome using two different strategies. In one case, we kept fixed the phenotype information and used gene dropping to generate genotypes for markers with allele frequencies corresponding to the one observed in our sample; in the other case, we kept the observed genotypes and permuted the phenotype values (with appropriate consideration of family structure). Considering all sets of simulations, HLODs of 4.1 or greater were seen in 80% of replicate genome screens, suggesting that, in these pedigrees, 4.1 is likely not an appropriate threshold for declaring genome-wide significant linkage.

**Figure 3.**
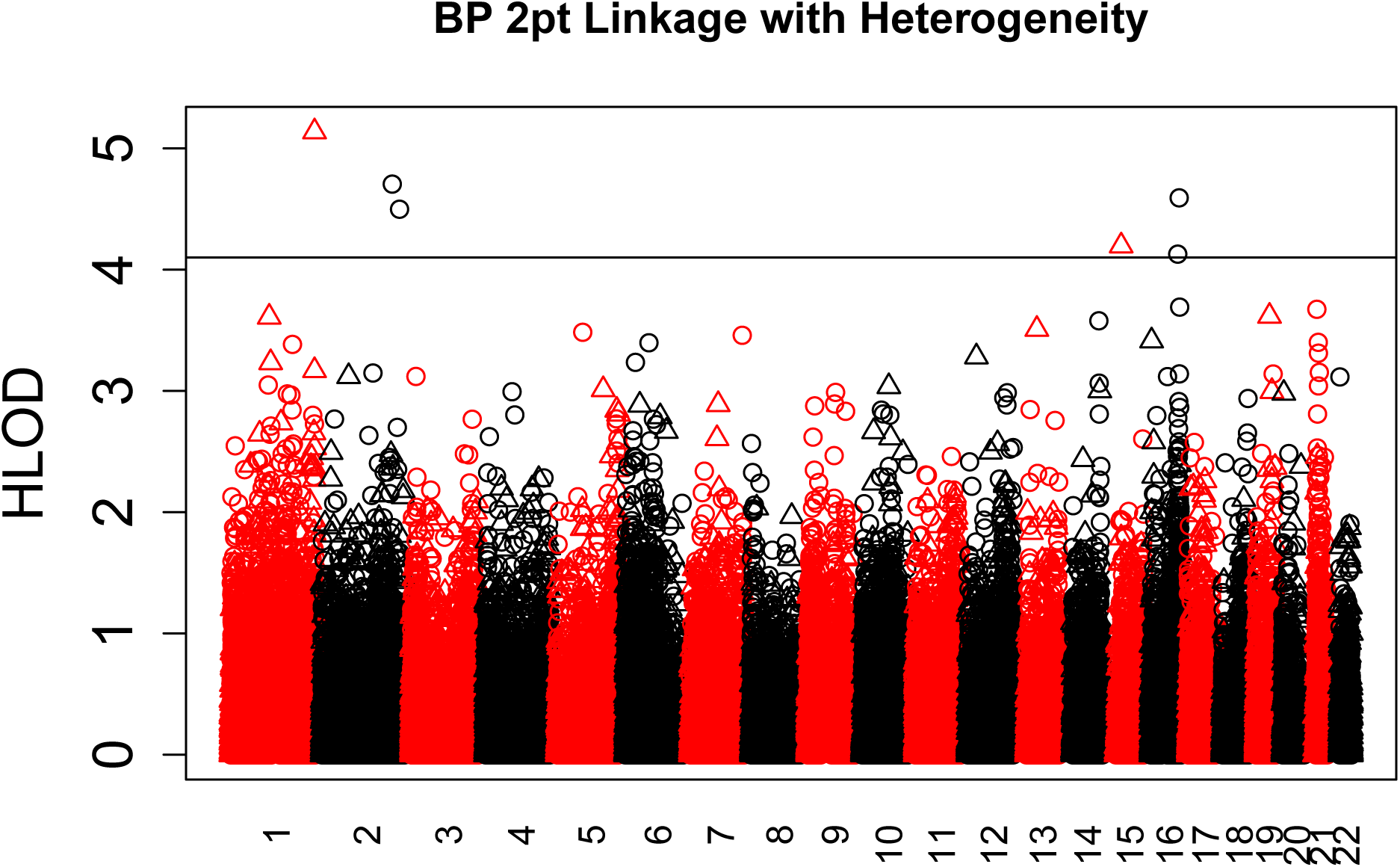
Manhattan plot of genome-wide 2pt parametric heterogeneity linkage analysis with 99,446 SNVs (circles) and 85,813 STRs (triangles). The horizontal line at HLOD=4.1 represents genome-wide significance.

**Table 3.**
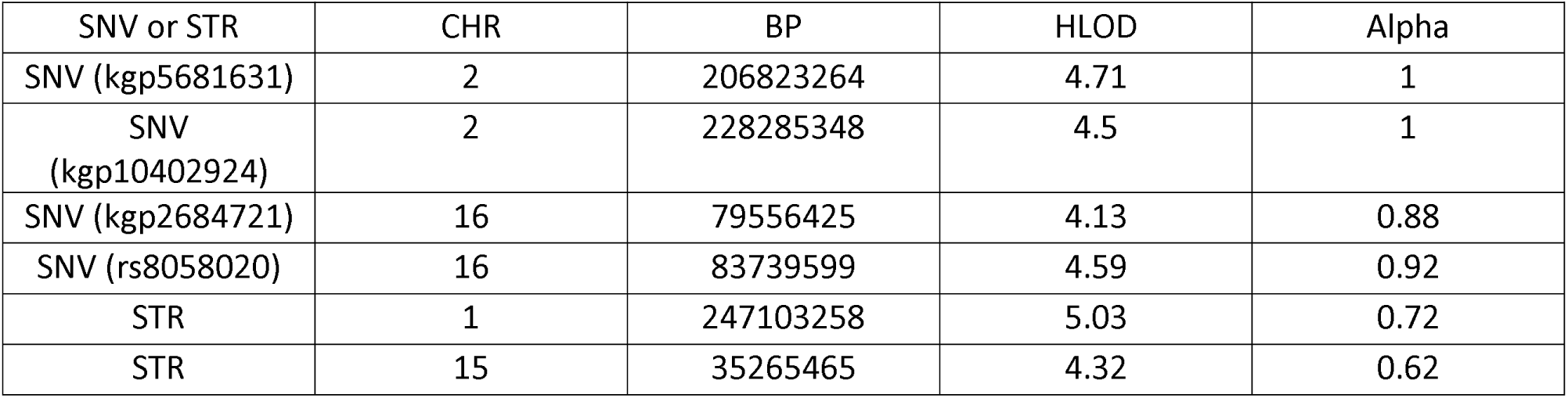
SNVs and STRs with heterogeneity LOD score (HLOD) to exceed 4.1. AN BP positions are on build 37, hgl9.

| SNV or STR | CHR | BP | HLOD | Alpha |
| --- | --- | --- | --- | --- |
| SNV (kgp5681631) | 2 | 206823264 | 4.71 | 1 |
| SNV<br>(kgp10402924) | 2 | 228285348 | 4.5 | 1 |
| SNV (kgp2684721) | 16 | 79556425 | 4.13 | 0.88 |
| SNV (rs8058020) | 16 | 83739599 | 4.59 | 0.92 |
| STR | 1 | 247103258 | 5.03 | 0.72 |
| STR | 15 | 35265465 | 4.32 | 0.62 |

In the NPL analysis, we approximated the empiric p-values using Rapid, and 13 SNPs and 156 STRs resulted in –log10(P)>3.5. These SNPs and STRs were re-analyzed with 250,000 simulations to obtain a true empiric p-value, and among those, eight SNPs and one STR had p<4.9e-05, which corresponds to the HLOD threshold > 4.1. We re-analyzed those eight SNPs and one STR with 1,000,000 simulations, and empiric p-values from this round of simulations ranged from 2e-06 to 3.9e-05, including two SNPs with p=2e-06, one on 16q23 (at 77.7 Mb) and one SNP on 21q21 (at 23.9 Mb) (Table 4).

**Table 4.** Nonparametric linkage results of 9 markers with empiric p-values <4.9e-05. Empiric p-values were initially tested wih 250,000 simulations and also with 1,000,000 simulations. BP Position is on build 37, hgl9.

| SNV or STR | CHR | BP | emp_pvalue<br>250,000<br>simulations | emp_pvalue<br>1,000,000<br>simulations |
| --- | --- | --- | --- | --- |
| SNV (rs804129) | 1 | 15183309 | 2.80E-05 | 2.80E-05 |
| SNV (rs3820546) | 1 | 43398085 | 3.20E-05 | 3.90E-05 |
| SNV<br>(kgp10316684) | 1 | 182297222 | 8.00E-06 | 1.10E-05 |
| SNV (rs10056131) | 5 | 28302858 | 1.20E-05 | 2.20E-05 |
| SNV (kgp9442190) | 6 | 31055539 | 2.80E-05 | 3.10E-05 |
| SNV (rs12444317) | 16 | 77705268 | 0 | 2.00E-06 |
| STR | 18 | 72256481 | 4.00E-05 | 2.60E-05 |
| SNV (kgp3613333) | 21 | 23959348 | 8.00E-06 | 2.00E-06 |
| SNV<br>(kgp10714796) | 21 | 24623957 | 1.60E-05 | 1.80E-05 |

#### Association analysis at the 16q23 locus

We sought to identify potential causal variants among the linkage peaks that we identified from SNPs and STRs. For this goal, we conducted association analysis on common variants that we detected in WGS data (190 BP1 and 130 controls). Among the 15 linkage peaks, we considered the 16q23 locus as the most likely linkage peak to contain identifiable causal variants, because of the linkage evidence observed with both parametric and non-parametric analyses. We calculated association statistics of 18 SNVs in the 16q23 locus that are putatively deleterious coding variants with a frequency > 5% in these pedigrees, 14 of which are located in a single gene, *PKD1L2*. None of these variants show significant association p-values after multiple testing correction (Table S8). We also found no difference in the mean burden of deleterious SNVs in *PKD1L2* between BP1 individuals compared with controls, controlling for differences in European ancestry (p=0.29 and 0.24 using LMM and GLMM, respectively).

#### Segregation of rare deleterious SNVs and CNVs in the gene-set for BP1

To detect the segregation of rare variants with BP1 in the CO/CR pedigrees, we developed a statistical approach that quantifies the significance of the observed segregation pattern. Intuitively, it estimates the probability that we would observe the given segregation pattern of a rare variant or more extreme patterns under the null hypothesis of random segregation; we refer to this as the segregation p-value. Our method used the imputation results to attempt to detect, with high confidence, founders who introduced a rare variant into a family. Given these founders, our method performed a gene dropping approach with random inheritance vectors (IVs) to find the proportion of IVs that generate the same or more extreme segregation pattern than the observed pattern, which becomes a p-value. For SNVs, we used the same set of 25,072 rare deleterious variants in our gene-set used for the burden analysis, while for CNVs, we used 922 rare bi-allelic deletions called from WGS data in 685 genes present in the gene-set after QC. We filtered variants not shared between BP1 individuals (Table S9). We were able to identify founders who introduced rare variants into the family with high confidence for 86.1% and 80.0% of all rare SNVs and CNVs, respectively (Figures S10 and S11); we kept all of these variants for segregation analysis. For 25.36% of SNVs and 35.20% of CNVs analyzed in the segregation analysis, more than one founder introduced rare alleles into a family. The fraction of such variants was greatest in the largest families (Figures S12 and S13). In total, we analyzed segregation for 6,421 rare SNVs in 4,050 genes and 314 rare CNVs in 251 genes.

No segregation p-value for either SNVs or CNVs was significant after correction for multiple testing (Tables S10 and S11). The top gene in the SNV segregation analysis was *ACTR1B* (p=5.18e-04). *ACTR1B* had one rare variant (rs141238033, chr2:98275876) present in two families (CR004 and CR201). In family CR004, the rare variant was introduced by a founder whose BP status was unknown, as was that of his children. Hence, for this variant we only analyzed segregation in family CR201. This variant was predicted by Polyphen-2 to be a damaging missense variant, did not appear in the Colombian samples within the 1000 Genomes data (MAF of 0%) and was very rare in the Latino samples of ExAC (MAF of 0.04%). It was enriched in the CO/CR pedigrees, with a MAF in all 449 sequenced individuals, not accounting for relatedness of 1.89% (47x over ExAC) and in family CR201 with 111 sequenced individuals of 6.76% (169x over ExAC). Accounting for relatedness(36) its MAF was 0.44% in CO/CR families (11x over ExAC) and 0.56% in family CR201 (14x over ExAC). *GOLPH3* was the top gene in the CNV segregation results (p=7.89e-4), with a single rare CNV (DEL_P0095_217, chr5:32161816-32162478) appearing only in family CO27. Its MAF among all sequenced individuals in the CO/CR pedigrees, not accounting for relatedness, was 0.78% and 14% in family CO27 with 25 sequenced individuals; accounting for relatedness its MAF was 0.22% in all families and 2.94% in family CO27. Neither of the above two rare variants segregated perfectly with BP1 status in the pedigrees in which they were present (Tables S12 and S13). In *GOLPH3,* we also observed a rare damaging SNV (rs192633198, chr5:32126511) in a different family (CO7) from the family carrying the rare CNV (CO27). The segregation p-value of this SNV in family CO7 was 0.12, but it segregated perfectly, having been inherited from a founder whose three offspring included two BP1 individuals carrying the rare allele and one unaffected individual who did not carry it.

### *De novo* mutations (DNMs) in CO/CR families

We discovered SNV DNMs among 67 trios using the WGS data with TrioDenovo software(22). After removing trios who carry zero DNM or more DNMs than expected, we identified 6,574 DNMs among 57 trios (115.3 DNMs per trio). We found no significant difference between BP1 and non-BP1 individuals in the burden of DNMs.

## DISCUSSION

The studies described here enabled us to document the contribution of both common and rare variants to BP1 risk, in the CO/CR pedigrees that we genotyped and sequenced. Our finding of elevated BP1 PRS scores, in BP1 affected individuals compared to controls, indicates that in these pedigrees, as in case/control samples, BP1 risk derives substantially from the polygenic effect of common SNPs. This result is consistent with what has been observed in extended pedigrees that apparently segregate non-psychiatric common disorders, such as familial dyslipidemias(37). It will, however, be important to understand why the magnitude of the polygenic contribution, in terms of the variance explained by PRS, is so much smaller in the CO/CR pedigrees compared to the cases from the PGC. The dissimilarity between the pedigree and case/control series in terms of sample size and ethnicity offers one explanation for this divergence(38). Another possibility is that our controls were closely related to BP1 individuals in our pedigrees, and hence they might carry some polygenic burden of BP1. Lastly, it is possible that BP1 in individuals from extended pedigrees is simply less polygenic than in population samples.

In support of the idea that BP1 in extended pedigrees may be etiologically distinct from this diagnosis in case/control samples, we found that common risk variants for SCZ do not appear to contribute to BP1 risk in our samples. This result contrasts with the observation in the PGC that the SCZ PRS is an equivalently strong predictor of BP1 risk as the BP PRS. This discrepancy relates to one of the most important uncertainties in our understanding of severe mental illness, the genetic relationship between BP and SCZ. The separation between BP and SCZ was for many decades a bedrock principle of psychiatric nosology, based on both the classically distinct longitudinal trajectories of these syndromes as well as evidence from genetic epidemiology studies suggesting that they do not co-segregate in families. More recently, however, several lines of evidence indicate a shared genetic architecture between SCZ and BP, particularly with respect to BP1(2, 25, 26). Efforts are now underway in multiple datasets to examine the relationship between BP and SCZ at a finer-grained level than that of syndromic diagnosis, for example in terms of the presence or absence of individual symptoms or the distribution of specific quantitative traits(39). Our results suggest that it may be particularly informative to include the rich phenotypic data available from the CO/CR pedigrees in such studies.

To date, there are no convincing reports of discoveries of high-impact BP susceptibility variants, or even loci, from either pedigree or case/control sequencing studies. Such studies have, almost certainly, all been underpowered for such discoveries. Many pedigree-based sequencing studies have, for example, described single families in which candidate variants were possibly segregating with disease(4-7). The dataset that we describe here is also underpowered for rare variant discovery. However, the comprehensive genotype data that it contains, for nearly 800 individuals from 22 families, provide an opportunity for more complete evaluation than has previously been possible of the contribution of rare variants to BP1 within pedigrees, and for a more rigorous examination of the segregation of rare variants with the disorder.

In assessing the contribution of rare variants to BP1, we found, in a set of 8,757 genes selected based on hypothesized relevance to this disorder, a higher burden of rare variants (deleterious SNVs as well as CNVs) in affected individuals compared to related controls. We chose this set of genes because they corresponded to regions where BP1 heritability was enriched and where BP1 GWAS hits and our linkage peaks resided. Ament et al. used a gene-set related to neuronal excitability and similarly observed enrichment of rare risk variants for BP among those genes(40). Previous studies have reported no evidence for a global enrichment of rare CNVs in BP individuals (41, 42). In contrast to these studies, however, which were restricted to analyses of CNVs > 100kb, our use of higher density arrays and WGS allowed us to examine much smaller events; our results strongly suggest that the impact of CNVs on BP burden derives mainly from smaller events (5-100kb), and may be restricted to specific sets of genes.

The discovery of increased rare-variant burden in the BP1 individuals led us to attempt to discover the specific loci and variants that increase risk for the disorder. Our linkage analysis yielded 15 peaks that harbor 99 genes. We analyzed in more detail the 16q23 locus, where, through parametric linkage we detected two SNPs with HLOD of parametric linkage > 4.1 and, through NPL a SNP with p < 4.9e-05. Analysis of 18 deleterious coding SNVs within this region yielded no associations to BP1, suggesting that non-coding variants may be responsible for the linkage result.

We applied, to this dataset, a new statistical approach that we developed to calculate a p-value for segregation of rare variants. We did so because existing methods used for this purpose (43-45) are not scalable to very large pedigrees such as those in our dataset. These methods also make the simplifying assumption that only one founder has introduced a given rare variant into the pedigree. While this assumption is reasonable for small or moderate-sized families, we have observed in large pedigrees (n>70) many instances in which more than one founder has introduced a rare variant into the family. Our method relies on accurate imputation of rare variants, which is made feasible by a family imputation approach that achieves a higher call rate and accuracy for rare alleles than population-based imputation approaches(17). This approach, however, requires reliable estimation of IVs in each family, and if only a small number of individuals in a family are genotyped at markers used for estimating IVs, imputation results might not be very accurate.

Although the method we developed worked well, from a technical standpoint, we did not detect rare variants with strong evidence of segregation in the CO/CR pedigrees. The top gene that we detected through SNV segregation analysis is *ACTR1B*. It has one damaging missense rare SNV that is very rare in the Latino population of ExAC, but has a relatively high MAF in one of our pedigrees, with the rare allele occurring at about 170X increased frequency compared to reference samples. A previous CNV analysis using WTCCC2 SCZ data found an association between the large duplication around *ACTR1B* and SCZ, but it was not replicated(46). *GOLPH3* is the top gene from the CNV segregation analysis, which contains one rare and relatively small deletion appearing only in one family. A previous linkage study for SCZ found a linkage peak on the 5p13 region that included this gene, but did not support association between *GOLPH3* and SCZ(47). A recent linkage study for BP also found a CNV duplication in GOLPH3 segregating with BP under the family-specific linkage peak(48).

One limitation in the segregation analysis for rare variants is that while we were able to identify founders who introduced rare variants to the family for a majority of variants, for some variants we could not identify those founders with high confidence, and hence did not test them. Duplications and CNVs from microarray data did not generate an allele for each chromosome, and hence they were not tested, as our method relies on imputation that requires genotype information. Lastly, we considered only deleterious coding SNVs for both burden and segregation analyses of rare variants; we did not analyze non-coding SNVs or indels because it is a challenge to measure their deleteriousness(49, 50).

In conclusion, our study demonstrates the polygenic genetic architecture of BP1 in a well characterized and large series of extended pedigrees, reflecting the action of a combination of many common and rare variants (including both SNVs and CNVs) with small or moderate effect sizes. Additionally, unlike in BP case/control samples, common SCZ risk alleles do not contribute to BP1 risk in these families. Finally, although our new method makes it feasible to rigorously evaluate rare variant segregation in large pedigrees, our inability to identify BP1-associated coding variants suggests that non-coding variants may play an important role in BP1 risk in these pedigrees.

## Methods

### Ascertainment and Phenotypic Assessment of Study Sample

The study sample consisted of members of 26 pedigrees, 15 from CR and 11 from CO, each ascertained based on multiple members who had previously been clinically diagnosed with BP1. Our group has studied three of the CR families(51, 52) and several of the CO families(53) in past linkage studies of BP1, however the composition of the families changed somewhat as we recruited new individuals. We increased the number of families for study by recruiting family members of BP1 individuals individually recruited for population-level mapping studies(54, 55). We expanded each family by systematically evaluating first degree relatives of the BP1 proband, and included other branches as we encountered individuals with suspected BP1. The ascertainment and phenotyping strategy employed in expansion of these pedigrees was previously reported in Fears *et al*.(13). Due to the way the pedigree was expanded, with an emphasis on collecting equivalent numbers of BP1 individuals and their unaffected relatives, and with systematic preferred expansion of branches with BP1 individuals, the BP1 phenotype itself is not heritable in this pedigree collection.

Diagnostic interviews were conducted using the Mini International Neuropsychiatric Interview (M.I.N.I)(56) and a Spanish version of the DIGS; the CO clinical team developed this translated and validated(57) version and made it publicly available at https://www.nimhgenetics.org/interviews/digs3.0b/. Interviews were performed by bilingual psychologists or psychiatrists extensively trained in the use of these instruments. Among CR individuals newly recruited for this study, only those who answered positively to M.I.N.I questions related to mood or psychotic symptoms were targeted for the complete DIGS interview. All available CO family members received a DIGS interview. Six research psychiatrists or psychologists, at the University of California, San Francisco (VR), University of California, Los Angeles (CEB, JGF, JM), University of Antioquia (CLJ), and Rutgers University (JE), who are experts in the diagnosis of mood disorders reviewed all available clinical material for the cases (DIGS interview, medical records, hospital notes), with at least one expert rater reviewing each case. Prior to beginning Best Estimate (BE) procedures each diagnostician established reliability (diagnostic agreement of 90% or better) with the Chair of the BE group (VR). Individuals designated as BP1 had a BE diagnosis of BP1, unipolar mania, or schizoaffective disorder, bipolar type. Control individuals were those who went through the complete psychiatric evaluation and were found to have no mental illness, as well as those who answered negatively to all M.I.N.I.(56) questions related to mood or psychotic symptoms, and were ≥60 years of age.

Written informed consent was obtained from all participants. Institutional Review Boards at participating institutions approved all study procedures.

### DNA Extraction and Microarray Genotyping QC

DNA was extracted from whole blood using standard protocols. Illumina Omni 2.5 chips were used for genotyping of 856 participants, in three batches. A subset of samples was repeated in each batch to enable concordance checks. A total of 2,026,257 SNPs were polymorphic and passed all QC procedures, including the evaluation of call rate, testing for Hardy Weinberg equilibrium, and Mendelian error. For linkage studies we used 99,446 SNPs with MAF>0.35, that had been LD-pruned to have r^2^<0.5. During QC procedures, allele frequency calculations, calculations of HWE, and estimates of LD were performed using only unrelated (founder) individuals. Further pedigree-wide Mendel checks were performed on the set of 99,446 SNPs used in linkage analysis; 0.7% of markers had 1 or more errors in Mendelian inheritance in these additional checks and all data for the marker in the family that generated the error was set to missing. Individual-level QC checks included verifying the pedigree structure by comparing theoretical kinship with empirical estimates, assessing missing rate, and verifying that the genetic sex agreed with the reported sex. Details of array genotype QC were published previously(11), and the final array genotype data set included data on 838 individuals in 26 families.

### Estimation of global admixture proportions

We generated estimates for all 838 pedigree members with SNP array genotype data. The reference populations were the CEU (n=112) and YRI (n=113) from HapMap(58, 59), as well as 52 Native American samples from Central or South America. The Native American samples were the Chibchan-speaking subset of those used in Reich et al.(60) selected to originate from geographical regions relevant to CR/CO and to have virtually no European or African admixture. In total the admixture analysis used 57,180 LD-pruned SNPs and 1,115 individuals. We compared the proportion of European ancestry between BP1 individuals and controls using both a linear mixed model (LMM) and a generalized linear mixed model (GLMM); both took into account relatedness of individuals using a kinship matrix. In LMM, the dependent variable was the burden score while the independent variable was BP1 status, and it was vice versa in GLMM. For LMM, we used the lmekin function in coxme R package(61), and for GLMM, we used the GMMAT software(62).

### Linkage Analysis

#### Overview

Several of the 26 pedigrees are large and complex, with multiple inbreeding or marriage loops. Such pedigrees pose a significant challenge to analysis, as standard multipoint approaches that employ the Lander-Green algorithm fail in this setting. Possible solutions would include trimming and cutting pedigrees into smaller units, however we found that applying this approach in these pedigrees led to a reduced capability to resolve phase, and therefore resulted in substantially reduced power. We also considered MCMC approaches(63), however given the density of markers in a genome-wide setting, such approaches are exceedingly slow, and we found it difficult to evaluate if we had reached convergence. We evaluated linkage in our pedigrees using a two-point parametric linkage on both SNPs and multi-allelic STRs identified from the WGS data using LobSTR(18), as well as a non-parametric method proposed specifically for use in large, complex pedigrees(35).

#### Two-point Parametric Linkage Analysis

We used the software Mendel(33) to estimate allele frequencies in the 26 pedigrees, performing these estimates separately for the CR and CO samples, and accounting for pedigree relationships. Two-point parametric linkage with heterogeneity was done in Mendel for autosomes, the very large pedigree CR201 had to be broken into four non-overlapping sections in order to do this analysis. Our parametric model used disease frequency to be 0.003, and penetrance parameters: P(BP1|DD)=0.9, P(BP1|DN)=0.81 and P(BP1|NN)=0.01; where “D” represents the disease allele and “N” represents the normal allele. Individuals are considered either affected with BP1 or phenotype unknown, that is, we designate no individuals as controls. This is the same model employed in McInnes et al.(51) and Herzberg et al.(53)

The null hypothesis of no linkage was evaluated using the LOD (logarithm of odds) score, which is the log 10 of the ratio of the likelihood of linkage, given the estimated recombination fraction, to the likelihood given the null value for the recombination fraction (0.5). Traditionally, a LOD score of 3 (p=0.0001) was considered significant linkage, however Lander & Kruglyak(34) suggested LOD=3.3 (p=4.9E-05) was a more appropriate threshold. The HLOD (LOD with heterogeneity), tests 2 parameters: the recombination fraction and proportion of linked pedigrees. The HLOD score has traditionally been evaluated using a combination of chi-square distributions with 1 and 2 degrees of freedom(64). By this metric, an HLOD of 3.6-3.7 corresponds to p^~^0.0001; using the Lander and Kruglyak p-value threshold of 4.9E-05 corresponds to an HLOD of 4.1.

#### Non-parametric Linkage Analysis

We used Rapid(35) for non-parametric analysis of allele-sharing of SNPs among BP1 individuals in our pedigrees. Rapid is designed for use in large complex pedigrees, where estimation of identity by descent (IBD) can be computationally demanding. Rapid evaluates significance using an approximation to the empiric p-value, which Abney et al.(35) states is conservative. After the first genome-wide scan to identify promising results using the approximation, we re-analyzed markers with –log10(P) > 3.5 with 250,000 simulations to obtain a true empiric p-value. Markers with p<4.9e-05 were then evaluated with 1,000,000 simulations to obtain a more accurate p-value.

#### Association analysis in 16q23 Locus

There are 18 common, deleterious SNVs from WGS data that reside in the coding regions of genes in the 16q23 locus identified by parametric linkage and NPL (between 77Mb and 84Mb). These are SNVs with MAF > 5% among founders, and we included SNVs that are stop-gain, stop-loss, splice-site, or missense variants predicted to be damaging by Polyphen-2(12). We computed an association p-value between BP1 status and each SNV using a generalized linear mixed model as implemented in the GMMAT software (62) GMMAT includes a kinship matrix as a variance component to account for relationships among individuals. We included the global admixture proportions of European ancestry as a fixed effect. Among the 18 SNVs, 14 SNVs are in *PKD1L2*, and we computed the mean burden of alternative alleles of these 14 SNVs using PLINK(65) ‐‐score option where missing genotypes are imputed. The comparison of the mean burden scores between BP1 individuals and controls was performed using both LMM and GLMM that take into account relatedness and global admixture proportions of European ancestry. In LMM, the dependent variable is the burden score while the independent variable is BP1 status, and it is vice versa in GLMM.

### Whole Genome Sequence (WGS) data

#### Individual Selection

We used ExomePicks (https://genome.sph.umich.edu/wiki/ExomePicks) to identify the subset of individuals to sequence that would enable maximum opportunity to impute sequence variants into the remaining genotyped individuals in the pedigrees. Selection in ExomePicks was done without regards to phenotype, when possible BP1 individuals were selected over non-BP siblings. ExomePicks selected 487 of the 838 genotyped individuals as candidates for sequencing. Sequencing was done in two rounds, and due to budgetary constraints, not all families were sequenced: no individuals from three CR families and one CO family were sequenced. In total 454 CR/CO individuals were sequenced, including 144 BP1 (Figure S1).

#### WGS and Variant Calling

Illumina performed WGS of 454 individuals using HiSeq 2000 with 36x overall coverage. Illumina performed initial variant calling using their internal variant caller, CASAVA, to obtain SNV calls in the VCF format. BAM files were also made available after the CASAVA variant calling pipeline. To improve accuracy of variant calls, we re-aligned and re-called the WGS data using the GATK best practices(14). We converted the original BAM files to FASTQ files, and used the Churchill pipeline(66), which is an efficient and scalable implementation of the GATK best practices pipeline. We used the HaplotypeCaller of GATK (version 3.5-0-g36282e4). First, each individual was called separately to generate a GVCF file, and all individuals were joint-called using the GenotypeGVCFs tool in GATK. The variant calling was performed in the high performance cluster at UCLA called the Hoffman2 cluster.

#### Individual-level QC of WGS Data

Before checking sequencing quality of each individual, we first removed variants that failed Variant Quality Score Recalibration (VQSR) in the GATK pipeline and we set each genotype whose genotype quality (GQ) score was ≤ 20 to missing. Additionally, all multi-allelic SNVs were excluded. In the remaining variants, for each sequenced individuals we assessed genotype missing rate, the number of singletons and WGS genotype concordance with array genotypes, verified agreement between reported sex and sex determined from X chromosome markers, and compared empirical estimates of kinship with theoretical estimates. We also performed principal component analysis (PCA) using 1000 Genomes (1KG) Phase 3 as a reference panel(16). We used EIGENSTRAT(67) for PCA, and included only founders from WGS data, and used independent common SNVs present in both BP1 WGS and 1KG Phase 3.

#### Variant-level QC of WGS Data

In doing variant-site QC, rather than using fixed thresholds for missingness, HWE, etc. to filter out poor quality variant sites, we used logistic regression to predict the probability of variant sites being of good or poor quality, with prediction based on mean and standard deviation of variant sequencing depth and genotype quality and a fraction of individuals meeting certain thresholds for depth and quality. We trained the regression model on a set of variants deemed to be of obvious good and poor quality using several QC measures such as genotype missing rate, HWE p-value, and the number of Mendelian errors. We assessed the accuracy of the model using cross validation (see Supplementary Text for details).

#### Genotype Refinement Using Pedigree-Aware Variant Calling Algorithm

We used the pedigree-aware variant calling method Polymutt(15) to refine genotypes of variant sites that passed QC. This method takes a VCF file with genotype likelihood generated from GATK and pedigree structure as input and generates a VCF file that refines genotype calls based on their likelihood and pedigree structure. To use Polymutt, some of large pedigrees had to be trimmed to remove inbreeding loops and reduce pedigree complexity. The use of Polymutt to refine genotype calls reduces the possibility of identifying *de novo* mutations, but results in a dataset with nearly zero Mendelian inheritance errors (MEs). Polymutt also imputed all sporadic missing genotypes.

#### Imputation of Sequenced Sites into Genotyped Individuals

The 454 individuals in 22 of the 26 CO/CR families who underwent WGS were selected with the goal of imputing the rest of 334 family members who were genotyped on the Omni 2.5 chip, but not directly sequenced (Figure S1). We used the imputation software GIGI(17), an approach designed to impute genotypes on large extended pedigrees. First, we obtained a genetic map of variant sites using the Rutgers genetic map interpolator(68). We then used MORGAN(69) to obtain pedigree IVs based on independent common SNP array genotype data of individuals in 22 families with WGS data. GIGI used these IVs to impute WGS variants identified in the directly sequenced individuals into the pedigree members with only SNP genotype data. GIGI also imputed sequence data for pedigree members who were neither sequenced nor genotyped in the Omni 2.5 chip. GIGI imputed each family separately, and we further divided each chromosome into 10 Mb intervals to perform efficient imputation by utilizing parallelization in the high performance cluster. After imputation, GIGI generated the probability of each imputed genotype and we used the threshold-based calling with the default threshold to call genotypes for the rare variant burden analysis while we used the most likely genotype calls for the rare variant segregation analysis. For the threshold-based calling approach, only genotypes with probability in excess of 80% were called, genotypes that did not meet this threshold were set to missing. After the GIGI imputation, there were 782 individuals who were either sequenced or imputed with the high quality in 22 pedigrees. Among them, 190 are BP1 and 130 are controls, and they were analyzed in analyses of polygenic risk, and burden and segregation for rare SNVs and CNVs.

#### Identification and QC of STRs

We detected STRs with the lobSTR software(18) (version 4.0.0) that uses sequencing data to call STRs. The BAM files aligned with BWA-MEM that were generated during the variant calling process were used as input to lobSTR, which then generated VCF files for STR loci. To perform QC on STR calls, we developed a filtering strategy based on the violation of Mendelian inheritance as well as comparison between lobSTR calls and STR data previously detected with electrophoresis for one family. This filter removed 1) monomorphic STRs, 2) STRs with repeats of one nucleotide, 3) STRs with ambiguous repeating nucleotides, 4) STRs with call rate < 95%, coverage < 5x, or Q-score < 0.95.

#### Identification of de novo mutations (DNMs)

There were 67 trios sequenced among 449 sequenced individuals after discarding five sequenced individuals with sequencing quality issues and possible sample mix-ups. We used TrioDenovo software(22) to identify DNM among the 67 trios with the minimum de novo quality (minDQ) of 8. We called DNMs on genetic variants that pass the GATK VQSR QC and on genotypes with GQ > 20. We removed variants present in dbSNP and variants whose minor allele count is > 1 among the 67 trios. We removed 10 trios that had zero DNM or more than 300 DNMs. DNMs were annotated with SnpEff software(70). We calculated association between BP1 status and the burden of DNMs using GLMM while taking into account relatedness.

#### Annotation of SNVs

Discovered SNVs were mapped to UCSC knownGene(71) and GENCODE V.19(72) transcripts, and damaging missense SNVs were predicted by PolyPhen-2(12). To identify rare SNVs, we used both external and internal sources of allele frequency. For external allele frequency information, we used allele frequency in 1000 Genomes(16) (1KG) Colombians (CLM) and ExAC(29) Latino population (AMR). If a variant in our dataset is present in 1KG CLM or ExAC AMR, it is rare if its MAF is < 1% in either dataset. If a variant is not present in both 1KG CLM and ExAC AMR, it is rare if its MAF is < 10% in our dataset where MAF is estimated from all sequenced individuals. We defined deleterious SNVs as variants that are stop-gain, stop-loss, splice-site, and missense variants predicted to be damaging by PolyPhen-2.

### Identification and annotation of rare CNVs

#### Array-based CNV detection

We adapted a previously established pipeline for array-based CNV detection (19); details are provided in the Supplementary Text. Briefly, after reclustering array data by genotyping batch, we generated a consensus callset based on two separate calling algorithms(73, 74). CNVs were only included in if they were called by both algorithms and of the same relative type (CNV gain or loss). We removed outliers based on several intensity-based metrics (Figure S4), merged fragmented CNV calls, removed CNVs in regions prone to false-positives, and further removed outliers based on global CNV load. Out of 838 samples passing SNP-based QC, 780 (93.1%) had intensity data suitable for CNV detection. We restricted our analysis to CNVs spanning at least 10 SNPs and > 5kb in length.

#### WGS-based CNV detection

We used Genome STRiP (20, 21) to both discover and genotype deletion CNVs across all samples with WGS. We removed three individuals with an excessive number of CNV calls who also failed SNV QC (Figure S14) and two individuals with possible sample mix-ups before genotyping deletions across all remaining individuals. Inspection of available trios (n=67) demonstrated low levels of Mendelian inconsistencies across these candidate sites (0.0026±0.00067, Figure S14). We filtered genotyped sites for redundancy and quality, generating a final WGS deletion callset consisting of 8,768 distinct CNV loci, and estimated a genome-wide FDR of 0.018 for deletions ≥ 5kb (Table S14). Finally, we removed monomorphic CNVs, Mendelian inconsistent genotypes, and genotypes with high missing rate > 5% prior to performing imputation of CNVs using the same imputation pipeline for SNVs. Details regarding CNV calling, filtering parameters, and post-processing steps are described in the Supplementary Text.

#### Annotation of rare CNVs

As with SNVs, we applied the similar method to classify rare CNVs in both array and WGS CNV datasets. We used two different sources of information, dependent on the platform, to exclude variants that are common in the general population. For array-based CNV calls, we extracted frequency information from Database of Genomic Variants (DGV) Gold Standard Variants (Release 2016-05-15)(30). Since the DGV is a consolidated resource comprised of many individual studies, we only considered frequency information for CNVs estimated from datasets with a minimum of 1000 distinct samples. For the WGS deletion call set, we used structural variant calls from Phase 3 of the 1000 Genomes Project(32), and extracted frequency information for all deletions in the AMR continental group. We annotated all CNVs in both the array-based and WGS datasets with frequencies from these public datasets based on a 50% reciprocal overlap, and retained CNVs with MAF < 1%. CNVs without frequency information were retained if their MAF is < 10% across all available samples in our dataset (regardless of phenotype).

### Identification of genes for burden and segregation analyses for rare variants

We highlighted genes for burden and segregation analysis of rare variants from three sources: (1) genes in regions that demonstrated evidence of enrichment of BP1 heritability, as evaluated in the PGC BP1 GWAS data set (summary GWAS statistics); (2) genes near PGC BP1 GWAS peaks; (3) genes within 1Mb of our linkage peaks. We only consider genes where there is at least one deleterious SNV in our BP1 dataset.

We identified regions with enrichment of BP1 heritability using PGC GWAS summary statistics and 220 epigenetic profiles that originated from 100 individual cell types or tissues. The epigenetic annotation came from cell-type specific histone modification (ChIP-seq) data generated by the NIH Roadmap Epigenome Project(75). H3K4me1, H3K4me3 and H3K9ac data were post-processed by Trynka et al.(76) Peaks were called using MACS v.1.4. For each cell-type and specific histone mark, start and end of the peaks were determined. H3K27ac data were post-processed separately by Hnisz et al.(77), also using MACS v.1.4. The 220 cell type-specific annotations were then divided into 10 groups by taking a union of the cell type-specific annotation within each group, following Finucane et al.(28) For each of the 10 tissue groups, the genome was annotated with the start and end of regions identified as active promotors/enhancers. Within all regions marked for a specific tissue group, we evaluated the possible enrichment of BP1 heritability using PGC BP1 GWAS summary statistics and Stratified LD Score Regression(28). For tissue groups that demonstrated significant BP1 heritability in the PGC, we then highlighted for further analysis in our BP1 families all genes within regions marked by the annotation process.

We also specifically targeted genes near genome-wide significant association signals in the PGC BP1 GWAS. The PGC BP1 GWAS summary statistics were clumped in PLINK, using our BP1 genotype data as the LD reference (founder genotypes only). Clumps were formed in windows of 250 Kb and using an r^2^ threshold of 0.1. Among the resultant clumps, if the lead SNV was genome-wide significantly associated to BP1 in the PGC data (p<5e-08), we determined the physical extent of the SNVs that were in the same clump as the lead SNP, and considered for further analysis all genes within such regions.

Lastly we targeted genes within 1 Mb of linkage peaks (both parametric and non-parametric) identified in our BP1 pedigrees. Linkage peaks were defined as parametric HLOD>4.1, or non-parametric p-value <4.9e-05.

### Polygenic Risk Score Analysis

We calculated the polygenic risk score (PRS) in our sample using summary statistics for autosomal SNVs from the PGC GWAS of BP1, and SCZ. SCZ GWAS results were downloaded from the PGC (https://www.med.unc.edu/pgc/results-and-downloads), and BP1 GWAS results were obtained with permission from the PGC. We used PRSice(78) with both sets of GWAS data. We excluded A/T and G/C SNVs and SNVs in the MHC region on chromosome 6. Our WGS data were LD clumped, and we retained from the GWAS summary statistics the most significant SNV for each clump. All PRS were estimated at five p-value thresholds, and we used GLMM to assess the odds of being BP1 as a function of increasing mean PRS, for each GWAS p-value threshold. We also used LMM, modeling PRS as a dependent variable and disease status as a predictor. Relationships among individuals were captured with a variance component to account for kinship(62), and global admixture proportions of European ancestry were included as a fixed effect in both LMM and GLMM. When estimating Nagelkerke R^2^, we used a logistic regression model without taking into account relationships as it was not straightforward to estimate R^2^ in the GLMM framework. We also did not include the global admixture proportions when calculating Nagelkerke R^2^ because we were interested in variance of BP1 explained only by the PRS.

### Comparison of the burden of deleterious SNVs and CNVs in BP1 and Controls

For each individual, we calculated the mean burden of rare deleterious SNVs using the PLINK ‐‐score option. This corresponds to the fraction of deleterious alternative minor alleles at those SNVs that each individual has. Missing genotypes were imputed by PLINK when computing the mean burden. We also calculated the mean burden of all rare SNVs in genes described as above including variants that are not deleterious. The mean burden of rare deleterious SNVs was regressed on the mean burden of all rare SNVs using LMM and included a kinship matrix to account for relatedness among individuals. The residuals of the mean burden of rare deleterious SNVs after LMM were then quantile-normal (QN) transformed. Similar to the burden analysis in the 16q23 locus, we compared the QN transformed residuals between BP1 individuals and controls using both LMM and GLMM while taking into account relationships among individuals and admixture proportions of European ancestry.

To test for an increased CNV burden in BP1 individuals compared to controls, we tabulated three measures for each individual: the total number of genes within our gene-set affected by CNVs (gene count), the total number of CNVs regardless of genic content (total CNV count), and the average size of all CNVs (average CNV size). Genes were considered affected by a particular CNV based on a strict overlap with annotated gene boundaries. To examine for global differences in CNV rate, we used a GLMM to compare total CNV counts between BP1 individuals and controls, accounting for individual relatedness and genotyping batch. For the CNV burden across our BP1 gene-set, similar to the mean burden of deleterious SNVs, gene count was first regressed on total CNV count and average CNV size while accounting for relatedness between individuals using LMM. The residuals were then QN transformed, and we used both LMM and GLMM to compare the transformed residuals between BP1 individuals and controls while accounting for relationships among individuals and global estimates of European ancestry. As gene count burden is influenced heavily by overall CNV detection rate (31), including total CNV count, average CNV size, and genotyping batch as covariates within the model is important to control for any case-control differences that may arise from variation in assay quality. Burden analysis for CNVs from WGS data was conducted in the same manner (without genotyping batch as a covariate). Although GenomeSTRiP genotypes individuals across all CNV loci with high efficiency, we note that inclusion of the total CNV count corrects for varying rates of imputation efficiency across samples.

### Segregation Analysis for rare SNVs and CNV

Given a rare variant in a family, we developed a statistical approach that computes a p-value to estimate the probability of having the observed segregation pattern or more an extreme segregation pattern. Our segregation statistic (S_rare_) is the sum of the number of affected (BP1) individuals with a rare variant and the number of unaffected (control) individuals without a rare variant. We assume that we know founders who introduced the rare variant into a family, and we denote these founders as F^rv^. We will discuss later how we can identify F^rv^ Given a S_rar_e statistic for a certain segregation pattern and F^rv^, we can compute a p-value of S_rare_ by enumerating random IVs assuming that F^rv^ have the variant and by finding the proportion of IVs that generate the same or larger S_rare_ values. We called this the “Family-level” p-value as we computed this p-value for each rare variant in each family. It is important to note that the same S_rare_ statistic in the same family may yield different family-level p-values depending on F^rv^. One example is illustrated in Figure S15 where there is a 5-generation family. In this family, among 10 individuals, only the latest offspring (individuals 9 and 10) are affected while all others have a missing phenotype, which generates the maximum S_rare_ statistic of 2. The p-value of S_rare_ statistic of 2 is much more significant when the top founder (founder 1) is F^rv^ compared to when a parent of individuals 9 and 10 (founder 8) is F^rv^. The fact that the rare variant is inherited in every generation and shared among the two affected individuals in the last generation is a more rare event than when the rare variant is directly inherited from a parent and hence yields a more significant p-value.

In addition to the family-level p-values, we computed a “Variant-level” p-value that meta-analyzed p-values across different families for the same rare variant. We combined p-values using the Fisher’s method (the sum of log p-values). The direction of effect was consistent across different families as we were only interested in the enrichment of rare variants among affected individuals and depletion among unaffected individuals, and we were not interested in the opposite direction. We also computed a “Gene-level” p-value that meta-analyzed p-values across different rare variants and families in a gene. We used Fisher’s method as well, and it is important to note that because of LD, a founder may have two or more rare variants in a gene with the same S_rare_ statistic and p-value. Because meta-analysis assumes independence among p-values, we included only one p-value from the same founder if there were multiple same p-values from this founder in a gene.

To detect F^rv^, founders who introduced the rare variant into a family, we used the results of GIGI imputation. GIGI generated the probability of imputed genotypes for everyone in a family, even those who were not genotyped with the microarray (including all founders). Assuming bi-allelic variants, GIGI generated three probabilities for three genotypes (1/1, 1/2, and 2/2 where 1 and 2 are two alleles). We had two strategies to identify F^rv^. In the first strategy, if the highest genotype probability among the three probabilities of a founder was > 0.8, we considered this founder to have a high-quality genotype. If all founders in a family had high-quality genotypes, we included this rare variant for analysis and identified F^rv^ who had a rare variant. We used the second strategy when all founders except a pair of top founders had high-quality genotypes. A pair of top founders was a couple who did not both have parents in the pedigree structure, and they were not usually sequenced or imputed well. In some cases, GIGI assigned about 0.5 probability of having a rare variant to both top founders. This indicated that a rare variant was inherited from one of the couples, but GIGI was not able to accurately identify which founder introduced the variant. Assuming that both top founders were neither sequenced nor imputed well, and they did not contribute to the S_rare_ statistic, the segregation p-value would be the same regardless of which of the founders carried the rare variant. Hence, we randomly assigned the rare variant to one of the top founders in the second strategy. We ignored rare variants in which there was more than one pair of top founder in a family who did not have the high-quality genotypes as well as rare variants in which founders carried two alleles of a rare variant. We analyzed rare variants with low genotype missing rate (< 5%) that at least two affected individuals shared in a family as we were not interested in rare variants present in one or zero affected individuals.

## Acknowledgments

We thank all study participants and thank the Psychiatric Genomics Consortium Bipolar Disorder group for sharing the latest BP1 GWAS summary statistics with us. This work was supported by NIMH grants R01 MH075007 and R01 MH095454, and NIEHS grant K01 ES028064.

## Figure Legends

Figure S1. An overview of steps to produce final QC’d array genotype and WGS data sets.

Figure S2. First two dimensions of a PCA analysis of founders from the BP1 WGS data set, using 1000 Genomes phase 3 samples as reference.

Figure S3. Comparison of genotype missing rate of individuals from imputed data and the genotype concordance rate between genotypes imputed by GIGI and microarray genotypes. Each point represents each imputed individual. Genotypes were missing if genotype probability from GIGI imputation was lower than the genotype probability threshold, and we included only individuals whose genotype missing rates were < 10% for subsequent analyses.

Figure S4. Array-based CNV detection sample QC. We excluded outlying samples based on manual inspection of the distributions across all samples on three different intensity-based QC metrics: (A) Log-R ratio standard deviation (LRR_SD), a general measure of intensity signal to noise; (B) B-allele frequency standard deviation (BAF_SD), the deviation of heterozygous B-allele frequency measures from the expect value of 0.5; (C) waviness (bottom), which measures the waviness of signal intensities after accounting for local GC-bias. (D) Additionally, we removed samples with an excessive number of rare CNV calls.

Figure S5. A de Finetti diagram showing global estimates of admixture proportions among African, European, and Native American ancestries in the CO/CR pedigrees. The global estimates were calculated using microarray data with ADMIXTURE software.

Figure S6. Comparison of global estimates of admixture proportions between BP1 individuals and controls. Admixture proportions were compared for African, European, and Native American ancestries separately.

Figure S7. Comparison of PRS estimated from PGC SCZ GWAS summary statistics in BP1 individuals and controls. PRS is computed at different GWAS p-value thresholds of the PGC SCZ GWAS.

Figure S8. Enrichment z-scores of heritability of BP in 10 different cell type groups using the stratified LD score regression. The horizontal dotted line is the threshold for significant enrichment. CNS: central nervous system. GI: Gastrointestinal.

Figure S9. Comparison of the burden of rare deleterious SNVs in BP1 individuals and controls in the 8,236 genes relevant to BP. The burden score was regressed on the burden of all rare variants in the 8,236 genes, and the residuals were quantile-normal transformed.

Figure S10. The number of rare SNVs passing the three different filters. The first filter (F1) is the number of rare variants with genotype missing rate < 5% and in which at least two affected individuals in a family share the variant. The second filter (F2) is the number of rare variants for which all founders have high-quality genotypes (the highest genotype probability among the three probabilities from GIGI imputation > 0.8). The third filter (F3) is the number of rare variants for which all founders except a pair of top founders have high-quality genotypes. The “Passing F1 & (F2 or F3)” filter is the number of rare variants that we analyzed in the segregation analysis.

Figure S11. The number of rare CNVs passing the three different filters. The first filter (F1) is the number of rare variants with genotype missing rate < 5% and in which at least two affected individuals in a family share the variant. The second filter (F2) is the number of rare variants for which all founders have high-quality genotypes (the highest genotype probability among the three probabilities from GIGI imputation > 0.8). The third filter (F3) is the number of rare variants for which all founders except a pair of top founders have high-quality genotypes. The “Passing F1 & (F2 or F3)” filter is the number of rare variants that we analyzed in the segregation analysis.

Figure S12. The number of rare SNVs that one, two, three, or four or more founders introduced into a family. These rare SNVs are SNVs for which we were able to identify founders with rare alleles confidently.

Figure S13. The number of rare CNVs that one, two, three, or four or more founders introduced into a family. These rare CNVs are CNVs for which we were able to identify founders with rare alleles confidently.

Figure S14. WGS CNV detection sample QC. (A) Following genome-wide detection of deletion CNVs from WGS, we removed five samples with an excessive number of CNV calls or with possible sample mix-up who also all failed SNV QC prior to imputation in additional samples with available genotype data (see Methods). (B) Per chromosome plot of total number of CNV calls detected by WGS across remaining samples reveals no substantial sample outliers. Peaks on chromosome 2 and chromosome 14 correspond to VDJ recombination regions. CNV calls within these regions were excluded from further analysis. (C) Rate of Mendelian-inconsistent CNV genotype calls across 67 complete trios available from all 449 sequenced individuals. The average rate of inconsistent sites across all genotyped sites was 0.0026±0.00067.

Figure S15. An example of a pedigree for the segregation analysis. Only individuals 9 and 10 are affected, and all other individuals have missing phenotypes.

